# Adaptive planning in human search

**DOI:** 10.1101/268938

**Authors:** Moritz J. F. Krusche, Eric Schulz, Arthur Guez, Maarten Speekenbrink

## Abstract

How do people plan ahead when searching for rewards? We investigate planning in a foraging task in which participants search for rewards on an infinite two-dimensional grid. Our results show that their search is best-described by a model which searches at least 3 steps ahead. Furthermore, participants do not seem to update their beliefs during planning, but rather treat their initial beliefs as given, a strategy similar to a heuristic called *root-sampling*. This planning algorithm corresponds well with participants’ behavior in test problems with restricted movement and varying degrees of information, outperforming more complex models. These results enrich our understanding of adaptive planning in complex environments.

## Introduction

An important trait of intelligent agents is their ability to plan ahead before performing actions, resulting in deliberate behavior that avoids costly mistakes. During planning, chains of hypothetical actions are played out until a certain depth based on what is known about the reward structure of the environment (Huys et al., 2015). For instance, De Groot (1978) found that amateur chess players can plan between 4 – 6 steps ahead when considering their next move. While flexible, planning is computationally expensive, and optimal solutions are frequently intractable (Dolan & Dayan, 2013). Recently, Balaguer et al. (2016) found that people can think ahead to solve complex problems by clustering steps into different contexts. Still, surprisingly little is known about how people plan ahead when searching for rewards. In part, this may be due to the inherent difficulty of separating exploration from planning.

We investigate participants’ search for rewards in a complex two-dimensional grid world. The task has a rich combinatorial structure in which the reward on a location depends on row- and column-parameters, and is particularly challenging for many approximate planning algorithms proposed in the machine learning literature. Our results indicate that participants learn in this task and improve their performance over time. Further computational model comparison shows that a search algorithm with a planning horizon of *at least 3 steps* describes participants’ decisions better than other cue-based or associative learning models. Importantly, this model uses a sampling heuristic which initializes beliefs once at the start and then reasons about what to do on subsequent steps, rather than also updating beliefs during the planning process. This provides a tractable solution to complex and dynamic planning scenarios. Our results advance our notion of how people plan ahead in a computationally challenging search task.

## Planning as tree-search

Tree-search algorithms are search algorithms that are used to plan ahead in complex reinforcement learning problems (Brown et al., 2012). Treating each state as a node of a tree, these algorithms plan ahead by expanding nodes via specific sampling routines. As exhaustive search of all possible action and state sequences is generally impossible, tree-search algorithms attempt to sample the most promising paths, ignoring improbable states and unfavorable actions. Because of their ability to solve challenging reinforcement learning problems, tree-search routines are highly popular in machine learning (Silver et al., 2016).

Several psychological studies have investigated the way people might perform tree-like planning before executing an action. In a relatively simple two-stage decision task, people show evidence of model-based planning, requiring them to think ahead for two steps in order to obtain maximum rewards (Daw et al., 2011).

In more complicated and longer sequential decision tasks, there is evidence that people adapt their search to aspects of the problem or task demands. For instance, Huys et al. (2012) showed that subjects adopted a simple tree-search strategy in which they curtailed any further evaluation of a sequence as soon as they encountered a large prospective loss (called pruning). Huys et al. (2015) used a model-based behavioral analysis to provide a detailed examination of participants’ performance in a moderately deep planning task and found that participants plan by establishing subgoals in a way that achieves a nearly maximal reduction in the cost of computing values of choices. Keramati et al. (2016) developed a three-stage decision task and found that increased time pressure led to shallower planning, suggesting that a speed-accuracy trade-off controls the depth of planning with deeper search leading to more accurate evaluation. Interestingly, their analysis revealed that subjects integrate habit-based cached values directly into goal-directed evaluations. Van Opheusden et al. (2017) also found that people perform shallower tree search under time pressure, but that they search more as they improve during learning. Taken together, these studies show that people can use clever heuristics to adapt the depth of their search by “pruning” relatively poor branches of the tree or using cached habit-based values to avoid further computation along a branch.

Here, we assess participants’ decisions in a challenging reward search task with (potentially) infinite states and depleting resources, providing challenging state-action-state dynamics.

## The potato farming task

To investigate planning in complex environments, we adapted a task first conceived as a challenging machine learning problem (Guez et al., 2013). In order to make it suitable for human participants, we framed the task as a foraging problem and implemented it as a browser-based game. Participants control a farmer on a grid-like field in pursuit of potatoes. Movement is possible horizontally or vertically using the 4 arrow keys (Figure 1). Every tile on the grid may yield a single potato when visited, and is thereafter depleted. The underlying structure of the task is determined by sampling, independently, for each row *i* a probability *p_i_* ~ Beta(α_1_, β_1_) and for each column *j* a probability *q_j_* ~ Beta(α_2_, β_2_). For a tile in row *i* and column *j*, the probability of success (a potato) is determined by the product of the two probabilities *p*(potato) = *p_i_* × *q_j_*.

**Figure 1:**
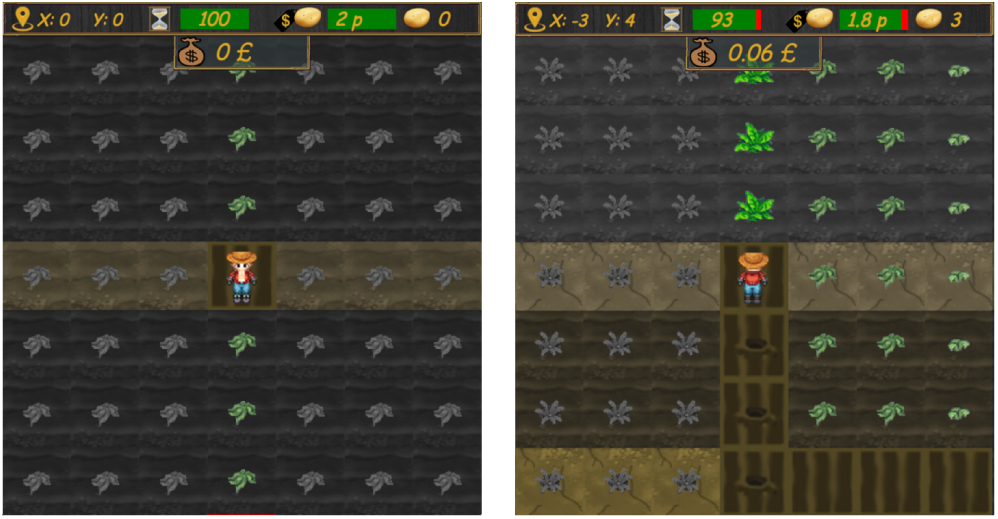
Screenshots of the potato farming game. The color and size of the plants as well as the color and texture of the soil provide visual cues for a tile’s quality. Further cues are delivered through the upper panel UI. The game can be played at https://git.io/vpSKp

While in theory infinite, the number of possible moves was limited to 100 (open) or 8 (test maps). Only a viewport of 7 × 7 tiles was shown during the game, with the farmer character on the center tile. Upon movement, both farmer and viewport shifted in the movement direction and the new center tile was harvested with the probability of success as stated above. Payoff values were —known to participants— temporally discounted with τ = 0.985 per move to further create opportunity costs for exploration, resulting in a 50% reduction of the original payoff per reward after 45 moves. The game contained distinct sound effects and animations that made reward pursuit more engaging.

In order to help participants infer the quality of rows and columns, tiles were displayed with visual cues. Different plant and soil types for each tile represented inferred column and row parameters, based on past reward frequencies (Figure 2). Specifically, there were 5 quality levels which were represented by distinct soil tones and textures, and by different plant shapes and sizes. Darker soil and larger plant types represented higher inferred quality. Additionally, the degree of information certainty, based on past exploration, was shown through overlaid shades of gray, with more gray representing less explored tiles.

**Figure 2:**
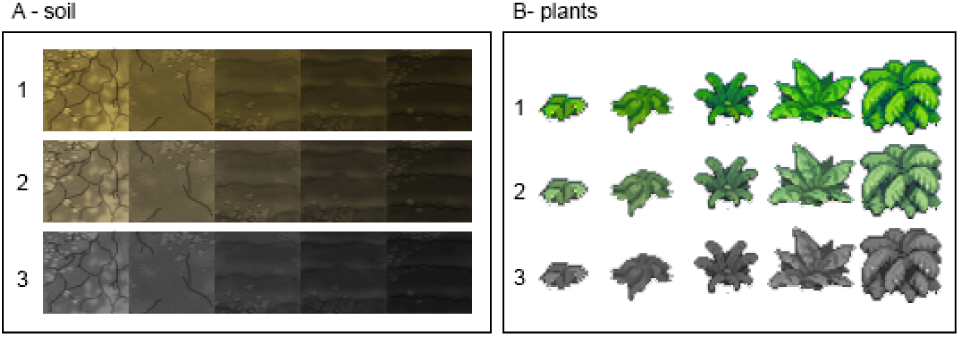
Visual cues based on past success and exploration. Displayed soil and plant types served as quality indicators for rows and columns (counterbalanced). The best soil/plant type can be seen on the right, the worst on the left. Shades of gray indicate information quality, with 3 being unexplored and 1 maximally explored.

Information certainty levels were determined by a monotonic rule based on the number of past moves within a row or column. The criteria were: high uncertainty (0 visits of row or column), medium uncertainty (between 1 and 4 visits), and low uncertainty (more than 4 visits). In contrast, inferred quality levels were presented based on a pseudo-Bayesian updating rule that took into account past exploration and success rates within a column or row. For every row or column *k*, this was calculated as

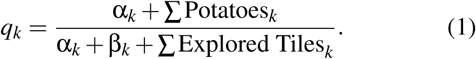

At the first move of every map, the player is naïve about the reward structure (Figure 1, left). After every subsequent move, the inferred quality and information levels for each visited row and column are updated (Figure 1, right). In order to maximize rewards, players must seek to find a route over the most rewarding tiles, for instance by traversing a column or row with a high parameter *p_i_* or *q_j_*.

The experiment was conducted online, and participants were recruited using the Prolific Academic platform (https://prolific.ac), a UK-based crowdsourcing platform solely focused on scientific studies. A total of 176 participants took part in the experiment, 8 of whom did not sent data, either due to technical problems (5) or for unknown reasons (3). Thus, 168 participants were included in the analysis (101 female, mean age=32.1). Participants were paid a flat participation fee of £2 and an additional bonus dependent on the number of potatoes harvested. The average reward over all participants was £2.51, and the average time taken was 19 minutes. Ethics approval was obtained from the UCL Research Ethics Committee.

The experiment contained 2 stages. First, participants navigated 5 open maps, each of which was dynamically generated and allowed for 100 unrestricted moves. Maps were created by sampling the row (*p_i_*) and column (*q_j_*) probabilities from one of two beta-distributions as a between-subjects condition. For one group the parameters were α_1_ = 1, β_1_ = 2, α_2_ = 2, and β_2_ = 1 (henceforth the *2-1-group*, Figure 3 right), and for the other group they were α_1_ = α_2_ = β_1_ = β_2_ = 0.5 (henceforth the *0.5-0.5-group*, Figure 3 left). This was done to create different reward structures, where rewards in the 2-1 condition were more clustered around rows or columns.

**Figure 3:**
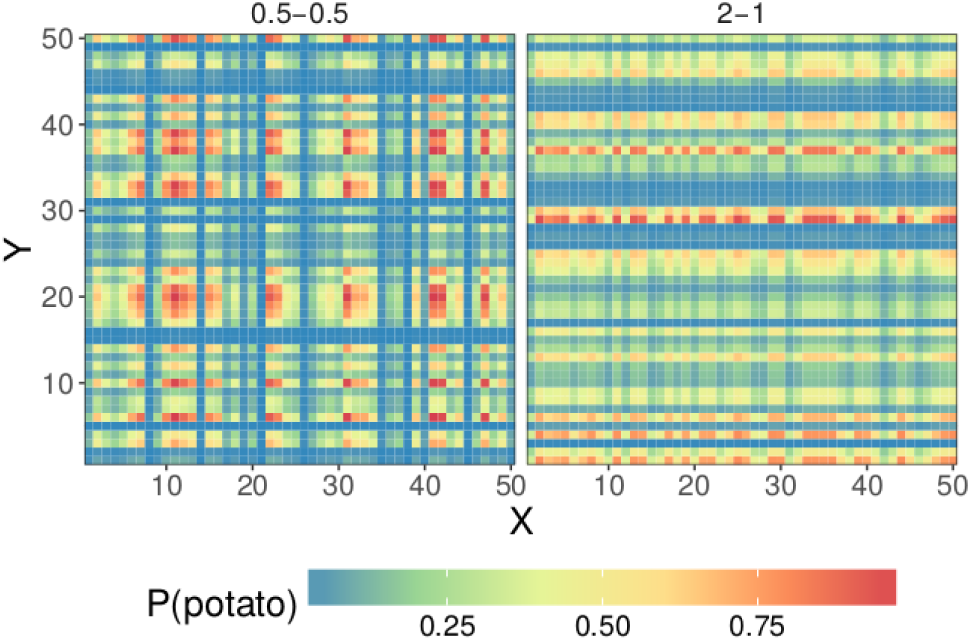
Between-subjects conditions for open maps. Probabilities were derived by multiplying realizations of two different Beta-distributions, i.e. the 2-1 and the 0.5-0.5 group. In the 2-1 group, following column *j* implies a collection of 2*q_j_*/3 reward on average (2/3 is the mean of a Beta(2,1)-distribution) whereas following any row *i* implies a collection of *p_i_*/*3* reward on average. In the 0.5-0.5 group, the row and column probabilities are more extreme and rewards more uniformly distributed over the map.

During the second stage, we presented 8 pre-designed test maps that included unnavigable tiles, displayed as water. The test maps were presented in random order. Every test map was created based on a simple trade-off between a proximal and almost certain reward and a superior, but distal option covering several tiles at the edge of the display. A limit of only 8 moves forced participants to pursue either option, and they could plan their moves from the start: In contrast to the open maps, information about both uncertainty and quality levels was revealed from the start (by using pseudo-counts, suggesting prior experience with columns and rows). There were two levels for both the degree of movement restriction and the simulated information certainty (see Figure 5). We hypothesized that higher certainty would lead to better planning and that more restrictions would lead to improved planning (as less options have to be considered).

## Behavioral results

### Open maps

We first analyzed performance in the open maps. Figure 4A shows participants’ average number of harvested potatoes over the 5 consecutive rounds of open map scenarios and Figure 4B shows the average rewards for the 100 trials (i.e., moves) per open map averaged over participants and rounds. To further assess the effect of trials, rounds, and condition, we regressed those variables onto rewards (i.e., whether or not a potato was gained) in a logistic regression. The results of this model are shown in Table 1.

**Figure 4:**
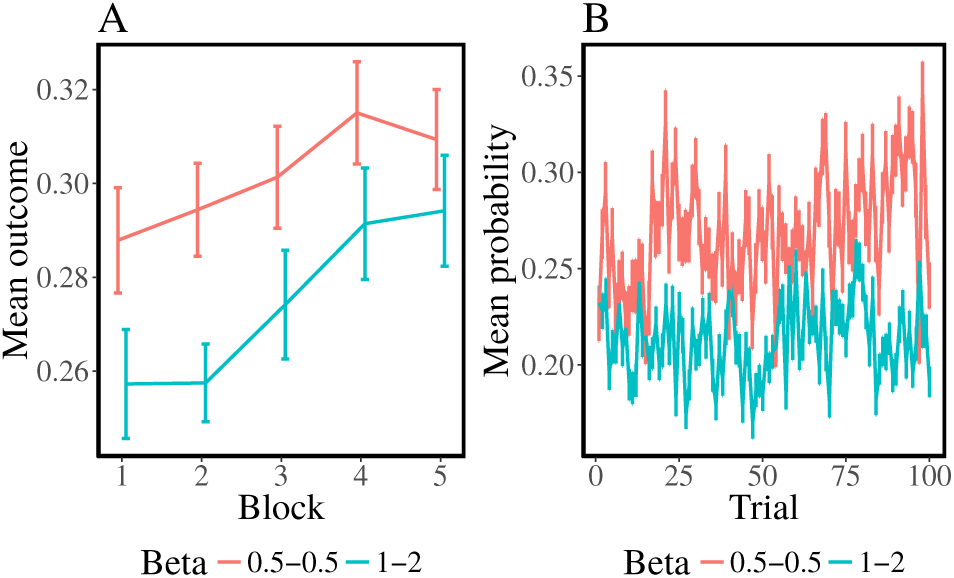
Behavioral results for open maps. **A**: Participants’ average reward per block. **B**: Participants average reward per trial.

**Table 1:**
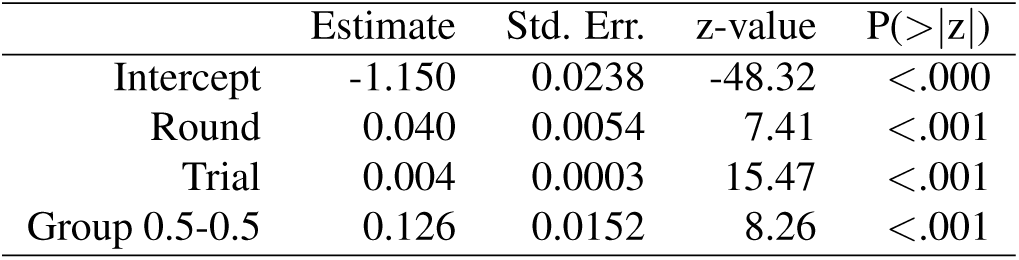
Results of logistic regression for behavior in open maps. Results are based on fixed effects estimates.

As expected, participants improved over rounds (β = 0.039, *p* < .001) and trials (β = 0.004, *p* < .001), indicating that they were able to learn in this task. Furthermore, participants in the 0.5-0.5 condition performed better on average than participants in the 2-1 condition. This means that performance is better if rewards are more extreme and evenly distributed rather than clustered within rows or columns.

### Test maps

Next, we analyzed participants’ performance in the test maps. Figure 5 shows a heat map of participants’ moves for each type of test map. Participant trajectories frequently aimed towards the distant, more promising tiles instead of the proximal but less promising ones, indicating that they were indeed planning ahead.

**Figure 5:**
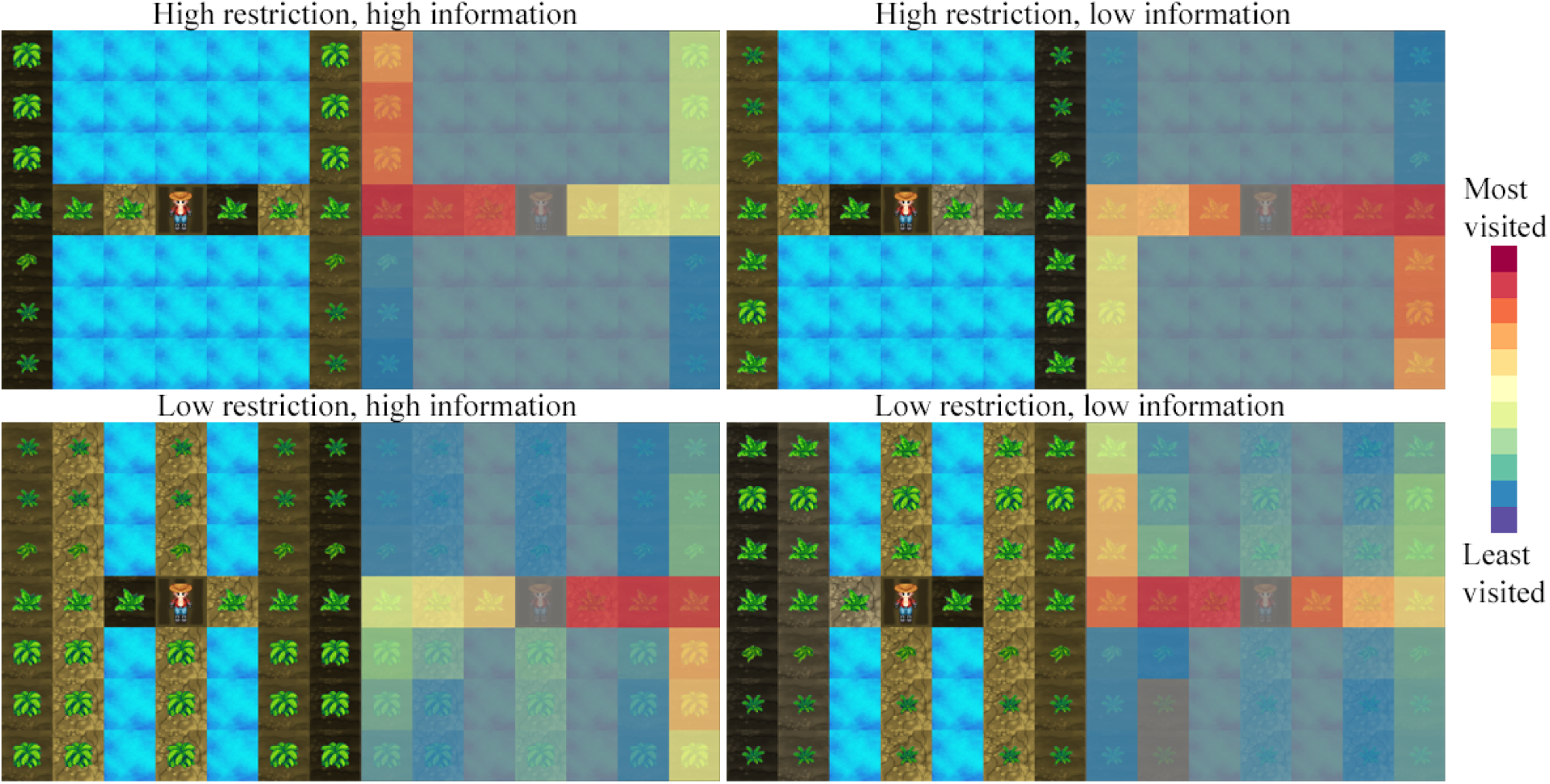
Test maps and corresponding heat maps. Two levels of movement restriction, visualized by water tiles, as well as two levels of information certainty, resulted in 4 combinations with distal rewards along one corner. Each combination also featured a 90 degree spatial rotation (not shown here) for a total of 8 maps. Adjacent heat maps highlight participant movement: The most common route always goes to the better distant location, indicating planning. Both more restriction of movements and more information lead to better planning.

To further assess participants’ behavior in the test maps, we created 2 dummy variables, one indicating high vs. low information and one indicating high vs. low movement restriction. Subsequently, we regressed these two variables onto participants’ total scores per test maps. Results of this regression (Table 2) indicate that participants performed better in test maps with higher information certainty. As can be seen in Figure 5, such maps feature more salient distal rewards. Even though the distal reward is always superior, increased salience might inhibit spontaneous movement and increase the likelihood of planning. Additionally, further restricting movement also improved performance. Thus, participants’ might be able to further focus their planning on promising paths if the number of options is limited.

**Table 2:** Results of mixed-effects regression for behavior in test maps. Shown estimates are simple fixed effects.

| | Estimate | Std. Err. | t value | $P(>\vert t\vert )$ |
| --- | --- | --- | --- | --- |
| Intercept | 3.79 | 0.076 | 49.83 | <2e-16 |
| High info | 0.19 | 0.087 | 2.24 | .02 |
| Restricted | 0.38 | 0.088 | 4.30 | 1.83e-05 |

## Model comparison

To further assess planning, we defined 4 candidate models that differed in how they plan ahead and generate predictions about a movement’s expected value, and assessed how well they describe participants’ moves on the open maps.

### Candidate models

The first model is a simple *Associative Learning model*. This model learns to select directions more frequently over time based on how often they have led to success in the past. It is completely feature-free and involves no planning whatsoever. Instead, each option’s estimated value at trial *t* + 1 is driven by an update of its valence based on the discrepancy between current valence and outcome, i.e. the prediction error:

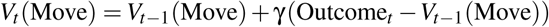

where γ is a free parameter governing the speed of the update over trials, i.e. how strongly the prediction error relates to adjustments of belief.

The second model is a *Neural Network model*. We fit this model to participants’ data from all but one blocks and then perform leave-one-block-out cross-validation. In particular, this model takes as input the current observable feature information in a game (the whole 7 × 7 matrix) at a time point *t* to predict a move at time point *t* + 1. To do so, we take a participant’s data for all but one block, find the best parameters to predict movements in this learning set, and then use the resulting neural network to make out-of-sample predictions for participant’s moves in the left-out block. This procedure is repeated for each participant and every block. The neural network consists of 1 hidden layer that can vary between a size of 1 to 4 nodes (best size chosen given all but the left-out blocks). The neural network model provides a good comparison to other, more explicit planning models as it takes in all available features at a time but is not based on any explicit planning (although it can capture planning implicitly).

The third model is a *Bayesian Monte Carlo search model*. This model takes in the current history of a participant’s moves and outcomes so far and then calculates the estimates of the row and column parameters (*p_i_* and *q_j_*) using the pseudo-Bayesian updating formula described in Equation 1. Afterwards, it plans ahead by executing random moves, collecting the sampled realizations of rewards from the multiplied probabilities, always performing a Bayesian update after each move and sampled outcome. Moreover, this model also correctly assigns a value of 0 to all previously visited tiles when planning future moves, thereby taking into account depleting resources. It is nonetheless still a relatively simplistic planning model as it does not better its moves as it looks ahead. We approximate the value of moves by using 10,000 random moves of a depth of *d* = {2,3,4,5} steps to assess how far participants plan ahead in our task. Although participants might plan further ahead than 5 steps, we did not include models of higher depth into our model comparison due to the computational complexity of assessing these models in our task (running a model of depth 5 for all participants currently takes around 2 days on a cluster with 64 active nodes). We use 10,000 random moves not because we assume that participants might actually mentally simulate that many trajectories, but rather to achieve an appropriate estimate for each move using Monte Carlo samples (i.e., our model would not work well with only a few samples due to the combinatorics of the task).

The final model is a *Heuristic Monte Carlo search model*. This models is similar to Bayesian Monte Carlo search, but instead of performing a Bayesian update after each simulated move, it only samples realizations of values for *p_i_* and *q_j_* once at the start and then executes all actions based on the initial samples. This strategy is related (but not equivalent) to an approach called *root sampling* as the probabilities are only sampled once at the root of the planning tree before the search commences (Guez et al., 2013). This heuristic can greatly simplify planning, especially in complex environments and therefore will be used as a heuristic planning model in our comparison. We again approximate the value of moves by using 10,000 random moves of a depth of *d* = {2,3,4,5} steps.

For all models, we use a softmax function to convert the value of an action *V_t_* (Move) into a choice probability

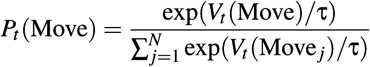

where τ is a free temperature parameter. As τ → 0 the highest-value action is chosen with a probability of 1, and when τ → ∞, all options are equally likely.

We calculate, for every participant and round, a model’s 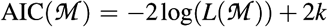 and use this to compute a pseudo-R^2^ measure as an indicator for goodness of fit, comparing each model 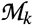 to a random model 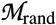:

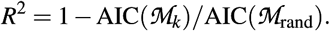

### Modeling results

Figure 6 shows the results of our model comparison procedure. The Heuristic Monte Carlo search models described participants’ moves better than both the neural network (*t*(167) = 2.89, *p* < .01, *d* = 0.22) and the Associative Learning model (*t*(167) = 23.31, *p* < .001, *d* = 1.80). The Bayesian Monte Carlo Search model also performed better than the Neural Network model (*t*(167) = 2.63, *p* < .01, *d* = 0.20) and the Associative Learning model (*t*(167) = 21.78, *p* < .001, *d* = 1.68). The Neural Network model described participants’ moves better than the Associative Learning model (*t*(167) = 7.38, *p* < .001, *d* = 0.57). Importantly, the Heuristic Monte Carlo search model performed better than the Bayesian Monte Carlo search model (*t*(167) = 4.72, *p* < .001 *d* = 0.36). Overall, 104 participants were best described by the heuristic Monte Carlo search model, whereas only 64 participants were best described by the Bayesian Monte Carlo search model. None of the participants was best described by either the neural network or the associative learning model. Further analyzing the heuristic Monte Carlo search model, a planning horizon of 3 described participants significantly better than a horizon of 2 (*t*(168) = 9.35, *p* < .001, *d* = 1.39), whereas planning horizons of higher than 3 did not lead to any further significant improvements (all *p* > .05). Thus, a Heuristic Monte Carlo search model with a planning horizon of a size of at least *d* = 3 provides a satisfactory description of human behavior in our task. However, we are currently not able to discriminate between planning horizons of more than 3 steps. Although one would normally conclude that given the same model class any AIC that is marginally better also implies better model fits, it is not always the case that selecting models this way automatically implements Occam’s razor such that often times functional parsimony should also be considered (Rasmussen & Ghahramani, 2001). Moreover, we do not have an appropriate way of accounting for the fact that deeper planning is computationally more expensive although all of these models have the same number of parameters; deeper planning always leads to a large increase of run times in our model comparison. Therefore, based on parsimony, we can currently only conclude that a planning horizon of *at least 3 steps* seems to be sufficient to explain participants’ behavior well in the open maps, but not exactly how many steps people plan ahead in our task. Table 3 shows the models’ overall AIC, trial-wise log-loss, and protected probability of exceedance, i.e. the likelihood that the proportion of participants captured by one model exceeds the proportion of participants captured by all other models (Stephan et al., 2009). This shows again that the Heuristic Monte Carlo model provides a comparatively good fit to participants’ data.

**Figure 6:**
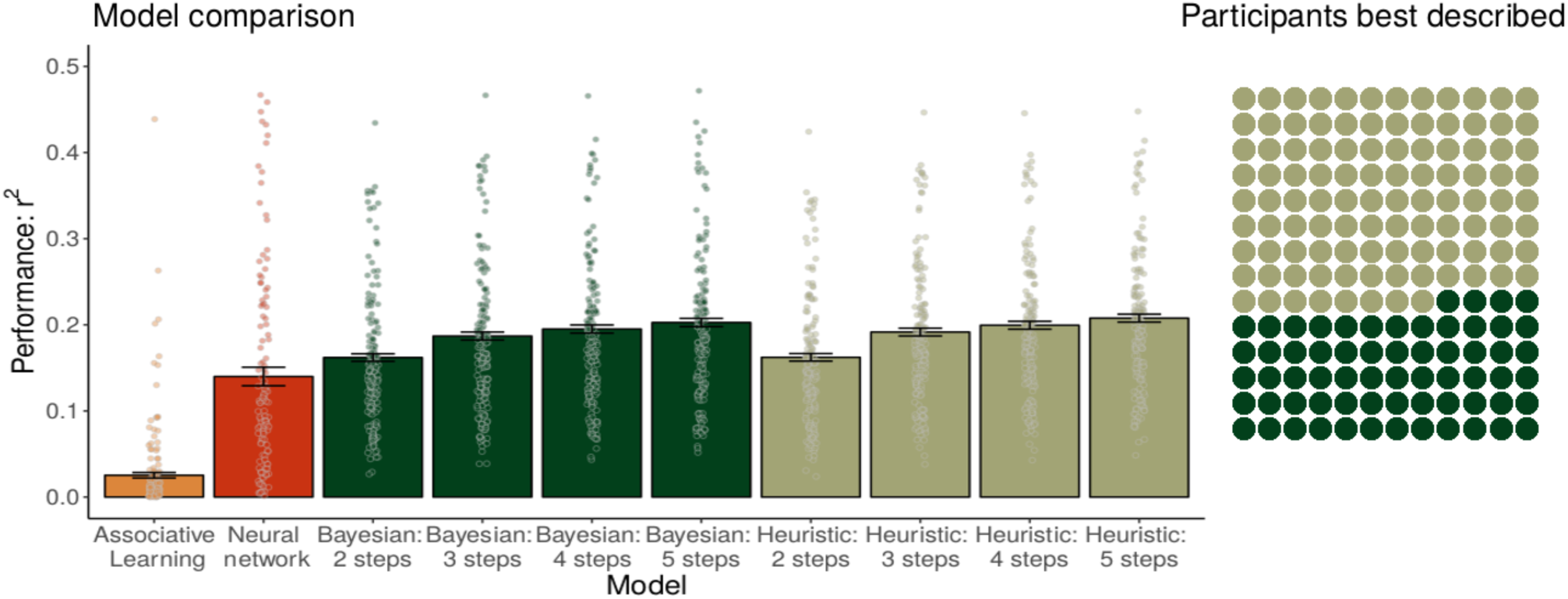
Results of the model comparison procedure. **Left**: Model fit for different models and horizon length. Error bars represent the standard error of the mean. Points show mean fit per participant. **Right**: Number of participants best described by each model.

**Table 3:** Model comparison results.

| Model | AIC | Loss | P. of ex. |
| --- | --- | --- | --- |
| Associative Learning | 1353 | -1.35 | .000 |
| Neural Network | 1194 | -1.19 | .005 |
| Bayesian Monte Carlo (3 steps) | 1107 | -1.11 | .010 |
| Heuristic Monte Carlo (3 steps) | 1100 | -1.09 | .985 |

Next, we assessed how well the different models performed at predicting participants’ moves in the test maps. For this, we used the parameters estimated from the open maps to perform out-of-sample predictions for participants’ decisions in the test maps, calculating the overall median performance of each model per participant over all test maps. As before, the Bayesian Monte Carlo model performed better than the Neural Network model (*t*(167) = 2.56, *p* < .05 *d* = 0.20) and the Associative Learning model (*t*(167) = 4.7, *p* < .001 *d* = 0.36). The Heuristic Monte Carlo model also led to better predictions for the test maps than the Neural Network (*t*(167) = 2.54, *p* < .05 *d* = 0.20) and associative learning model (*t*(167) = 3.55, *p* < .001 *d* = 0.27). As before, the Neural Network lead to better predictions for the test maps than the Associative Learning model (*t*(167) = 3.55, *p* < .001 *d* = 0.27). The Heuristic Monte Carlo search model captured participants’ behavior equally well as the Bayesian Monte Carlo search model (*t*(167) = 1.87, *p* = .07, *d* = 0.19). A horizon of 3 steps again turned out to be sufficient to explain participants’ behavior, describing participants’ decisions better than a horizon of 2 (*t*(167) = 2.78, *p* < .001 *d* = 0.21) and was again indistinguishable from a horizon of 4 or 5 (all *p* > .05). Taken together, these results imply that a simple model without belief updating, which plans the next move by sampling random sequences of moves with a horizon of at least 3 steps, describes human behavior in our task better than a purely associative reinforcement learning model, a simple neuronal network model, and other models of search with smaller horizons or more sophisticated belief updating.

## Discussion and conclusion

Planning ahead to search for rewards is an ubiquitous part of our everyday lives. Yet the process by which people do so has previously not gained much scientific attention. We investigated participants’ planning in a challenging foraging task with depleting rewards. Our results revealed that participants managed to learn in this task, improving their performance over time and rounds. Using computational modeling, we defined several models of planning and search and compared them based on how well they described participants’ moves in our experiment. This comparison revealed that people are best-described by a simple heuristic model of search that —instead of updating beliefs on every step— initializes beliefs only at the start and plans ahead for *at least* the next 3 moves. As this mechanism of planning has been found to lead to competitive performance and similar strategies can lead to optimal behavior (Guez et al., 2013), people seem to plan ahead heuristically yet efficiently in this complex search task. In future studies, we aim to assess the importance of generalization in a modification of this task in which outcomes are spatially correlated (Wu et al., 2017), compare more sophisticated mechanisms of exploration (Schulz et al., 2017a) as well as probe how introducing losses might influence participants’ search (cf. Huys et al., 2015; Schulz et al., 2017b).

Moreover, we intend to tease apart the different models of planning further by creating test sets to optimally discriminate between the models. We have also developed an updated version of the game that makes the visual aids even more explicit (see https://git.io/vpSPf). Importantly, our models only involved very simplistic planning by random roll-out; comparing more sophisticated tree-search models is therefore the most promising next step ahead.

## Acknowledgements

MK and ES contributed equally. We thank Peter Dayan and the Max Planck Research Group iSearch for helpful comments. MK is supported by the ESRC and Warwick Business School. ES is supported by the Harvard Data Science Initiative.

